# Efficient coding characterizes altered neural representations elicited by subtle sensory lesions

**DOI:** 10.64898/2026.04.10.717653

**Authors:** Juan Andrés M. Fuentes, Jaime Undurraga, Roland Schaette, David McAlpine

**Affiliations:** Center for Brain and Cognition, Universitat Pompeu Fabra, Barcelona, Spain; Australian Hearing Hub, Macquarie University, Sydney, Australia; Department of Linguistics, Macquarie University, Sydney, Australia; Interacoustics Research Unit, Technical University of Denmark, Lyngby, Denmark; Ear Institute, University College London, London, UK

**Keywords:** efficient coding, hidden hearing loss, cochlear synaptopathy, auditory system, neural adaptation, inferior colliculus

## Abstract

Sensory systems must represent a vast range of stimulus dimensions and energy whilst subject to metabolic constraints. Efficient-coding theory predicts that neural adaptation re-allocates a relatively limited range of neural activity toward the most informative stimulus values, but it is unclear how subtle peripheral lesions shift this operating point in central circuits. Hearing is a stringent test because sound level varies enormously across environments, yet clinical assessment still relies heavily on tone-detection thresholds that can miss listening deficits in noise. We analyzed extracellular recordings from single neurons in the gerbil auditory midbrain across 14 animals in four experimental groups exposed to unfolding distributions of sound intensities drawn either uniformly from a wide range (24-96 decibels) of sound pressure levels or from contexts in which 80% of levels were restricted to a 12-decibel high-probability range. For each context we summarized each neuron’s rate-intensity input-output function by an effective threshold and gain, and we interpreted the resulting threshold-gain distributions with an information-cost model that trades bits of stimulus information against a penalty on mean spiking. Noise exposure consistent with loss of synapses between inner-ear cells and auditory nerve fibers altered gain modulation across acoustic contexts, with noise-exposed animals showing compressed gain adjustments relative to controls; within the information-cost framework, the clearest hidden-hearing-loss effect was a quiet-context utility advantage distributed across low- and intermediate-threshold neurons, whereas moderate-to-loud contexts showed weaker or absent group differences. Temporary conductive attenuation caused by ear-canal plugging shifted effective thresholds to higher sound levels, with incomplete recovery after plug removal; the corresponding optimization-prior trajectories were consistent with incomplete rapid renormalization but were weaker than the hidden-hearing-loss effect. These results support an efficient-coding interpretation of altered central auditory representations after subtle lesions and provide a quantitative, context-based framework for comparing mechanisms of hearing difficulty beyond threshold-only tests and Fisher information alone.

**Author Summary:** Everyday hearing is an ecological challenge for the auditory system: we must follow speech while background sounds fluctuate and overlap. Standard tests emphasize tone-detection thresholds, but many listeners struggle in noise even when thresholds appear normal. We asked whether subtle peripheral changes shift how the auditory brain trades information for neural effort. We analyzed recordings from single neurons in the gerbil auditory midbrain during sound environments with different loudness statistics, including ones dominated by a narrow intensity range. Using information-theoretic measures, we quantified how much spikes distinguished sound-level categories and related this to the amount of spiking produced. Noise exposure consistent with inner-ear synaptic loss altered gain modulation across acoustic contexts and most strongly improved model-based coding utility in quieter settings, but reduced adaptation and efficiency as sound environments became louder. Temporary ear-canal plugging raised effective response thresholds substantially above both control and synaptopathy groups, with only partial recovery immediately after plug removal. By mapping both manipulations onto a common information-versus-cost scale, we highlight context-dependent metrics that may prove more informative than threshold audiograms for subtle hearing problems.

## Introduction

Sensory systems must extract and process (‘make sense of’) a wide range of stimulus dimensions and ranges of energy within those dimensions; sound intensities, for example, can span twelve orders of magnitude, from the softest sounds perceptible in a quiet room to a nearby jet-engine during take-off. Given resource constraints, it is likely that the brain represents this high dimensionality in an efficient manner. One way it achieves this is through adaptation (Brenner et al., 2000; Fairhall et al., 2001), the process by which response properties of individual and populations of neurons change to better reflect the current (over some defined epoch) statistical properties of the sensory environment (Carandini and Heeger, 2012). Adaptation takes advantage of the fact that whilst the long-term dynamic range of sensory input might be broad, this is not necessarily the case over shorter timescales. Conversational speech, for example, typically spans a range of 12 or so decibels in any setting, rather than the full 120 dB available to the human ear (Moore et al., 2008; Wen et al., 2009). Adapting to the unfolding statistical structure of sensory environments can reduce the representation of stimulus range to tractable dimensions (Willmore and King, 2023), and contributes to behavioral sensitivity to physical units over many orders of magnitude (Dean et al., 2005; Klotz-Weigand and Enz, 2022). Here, we explore whether altered neural representations of sound environments in the auditory midbrain (inferior colliculus, IC) following subtle peripheral lesions can be explained in terms of changes in coding efficiency, specifically a trade-off between the transmission of information and metabolic requirements.

The efficient coding hypothesis (Barlow, 1961), is key to understanding the role of adaptation in sensory representations; it argues that adaptation maximizes the information that sensory systems convey about the environment (Brenner et al., 2000; Van Hateren and Ruderman, 1998; Wark et al., 2007; Weber et al., 2019). Quantitatively, efficient coding is implemented as an optimization problem. It maximizes statistical quantities (such as mutual information, a measure of the mutual dependence between two variables) whilst imposing metabolic constraints, expressed through regularization terms representing mean neural firing-rate (Młynarski and Hermundstad, 2018; Gjorgjieva et al., 2019) and/or the sparsity with which neural firing patterns are generated (Olshausen and Field, 1996; Młynarski and Hermundstad, 2021).

Efficient-coding models suggest that the placement and spacing of neural thresholds (i.e., the sigmoid midpoints that determine each neuron’s operating range) and their diversity across a population can be derived from information-theoretic objectives under metabolic constraints, yielding specific predictions for how thresholds should partition stimulus space and how this partitioning depends on noise (both intrinsic variability within neural circuits and external sensory noise) and coding goals (Gjorgjieva et al., 2019). Related, the optimal structure for neural thresholds depends strongly on the specific location at which noise arises within (or is introduced to) a sensory pathway and on whether neural activity is combined across channels or preserved as independent channels. These features generate qualitative changes in the distribution of optimal thresholds as effective noise increases (Röth et al., 2021). Crucially, all such predictions assume a specific end goal (typically, maximizing mutual information between stimulus and response under a constraint on mean firing rate or total metabolic expenditure), so that changes in the constraint or in the noise statistics can shift the optimal code to a qualitatively different regime.

Here, rather than asking simply what should be the optimum distribution given a fixed task and noise model, we treat two forms of subtle lesions to the peripheral auditory system as perturbations that shift the inferred operating point of auditory neurons within an information-cost landscape. Classically, this framework has been applied to sensory peripheries, where it explains receptive-field structure and adaptation in vision and audition, but recent work extends the same information-metabolism trade-offs to central circuits and perceptual behavior, suggesting that efficient coding may provide a unifying principle across sensory modalities and processing levels. The efficient coding hypothesis and its mathematical implementation have the potential to explain the structure and activity of sensory end-organs and nervous systems, and to characterize ecological and evolutionary relationships, between the properties of, for example, cochlear filters and complex sounds such as speech (Smith and Lewicki, 2006), or the characteristics of retinal receptive fields relative to the features comprising natural images (Lörincz et al., 2012; Młynarski et al., 2021). The efficient coding hypothesis has also been used successfully to explain response properties of sensory neurons in terms of optimization related to information-theoretic quantities (Atick and Redlich, 1990; Atick, 1992; Van Hateren, 1993; Smirnakis et al., 1997; Smith and Lewicki, 2006; Dean et al., 2005, 2008; Watkins and Barbour, 2008; Willmore and King, 2023; Młynarski and Hermundstad, 2018). Nevertheless, there are no studies of which we are aware that have employed principles of efficiency to assess the consequences of deficits in sensory processing on the functioning nervous system (though see (Eckmann et al., 2020) for an explanation of the development of binocular vision, and how amblyopia can be considered a degeneration in coding efficiency). Assessing sensory deficits through phenomenological models of information transmission and efficiency has the potential to extend the understanding of sensory systems to cases in which they underperform given biophysical changes arising from perturbations of, or damage to, sensory end organs, with the goal of associating behavioral outputs to diminished efficiency of sensory representation. Sensorineural deficits might arise from biophysical changes at any stage in the afferent and/or efferent pathways, modifying their normal adaptive operation (McGill et al., 2022), and potentially diminishing the efficiency of an adaptive neural code in representing statistical properties of dynamic environments. Nevertheless, a challenge to understanding the consequences of sensory impairment on the organization and function of the central nervous system comes from the relatively uncontrolled consequences they elicit.

Damage to sensory end-organs can generate a cascade of changes in brain structure, connectivity, and excitability (Resnik and Polley, 2021; McGill et al., 2022; Chambers et al., 2016), and although maladaptive outcomes, such as phantom limb pain, potentially represent the consequence of homeostatic mechanisms in the central nervous system to retrieve a state of optimality given altered input, the potentially extensive and permanent damage to sensory surfaces with concomitant changes in brain function likely make it difficult to explore subtle changes in neural coding arising from principles of coding efficiency acting on altered sensory input.

One specific form of sensory deficit for which such an approach is feasible, however, and one that has attracted much attention in the last decade, is ‘hidden hearing loss’ (HHL), a colloquial term coined to encompass hearing difficulties in noisy environments without changes in sensitivity (thresholds) of the type normally associated with deafness (Schaette and McAlpine, 2011; Plack et al., 2014). A key underlying pathology contributing to HHL is cochlear synaptopathy, damage to a specific population of auditory nerve fibers (ANFs) in the inner ear, the high-threshold fibers considered important for coding louder sounds in background noise and potentially contributing to system gain (Bharadwaj et al., 2014; Hickox and Liberman, 2014; Bakay et al., 2018). Exposure to a single episode of 2 hours of 100 dB SPL noise in rodents leaves outer and inner sensory hair cells (and hearing thresholds) intact but reduces permanently supra-threshold neural activity in the early auditory pathways(Kujawa and Liberman, 2009). Cochlear synaptopathy is proposed to manifest as commonly reported hearing problems such as tinnitus (Schaette and Kempter, 2006; Schaette and McAlpine, 2011), hyperacusis (Hickox and Liberman, 2014; Zeng, 2013), and difficulties listening in ‘cocktail party’ environments (Auerbach and Gritton, 2022). To date, most *in vivo* studies of HHL have focused on understanding the consequences of noise exposure on ANFs, including the extent to which some of these changes might be reversible. Nevertheless, several studies have explored the consequences of HHL on central auditory processing, including the neural representation of speech-in-noise and adaptive processes contributing to dynamic range coding. Most recently, IC recordings in noise-exposed rats with confirmed synaptopathy have characterized changes in frequency, intensity, and duration coding consistent with reduced afferent sampling of the acoustic scene (Bakay et al., 2024), complementing the present study’s focus on adaptation to sound-level distributions in the same midbrain station. Sensory adaptation extends the range of hearing by allocating more neural coding resources to regions (of sound intensity) that appear with high probability (Dean et al., 2005, 2008; Auerbach and Gritton, 2022). In animals with suspected HHL, sensory adaptation to unfolding distributions of sound intensities is disrupted, particularly for louder contexts (Bakay et al., 2018). Paradoxically, for sound distributions with average low and moderate intensities, coding capacity appears better in HHL animals than in controls, consistent with the neural representation of speech in background noise being better at moderate compared to higher overall sound intensities for the same signal-to-noise ratio (Monaghan et al., 2020).

A still more subtle form of sensory impairment than HHL, and one potentially completely reversible, is conductive hearing-loss (CHL): physical changes in the outer and middle-ear that attenuate the transmission of sound energy to the inner ear, ultimately elevating hearing thresholds (Sataloff and Roehm, 2024). CHL can arise from relatively simple blockage of the ear canal by the buildup of earwax or insertion of foreign bodies or, common in childhood, bacterial or viral infection of the middle ear generating viscous fluid (‘glue ear’) behind the ear drum (Lloyd et al., 2021; Aithal et al., 2012). Long-term, CHL can generate comparable impacts to nerve degeneration or synaptopathy (Okada et al., 2020; Liberman et al., 2015; Manno et al., 2021; Thornton et al., 2021). CHL can be induced experimentally in vivo or in humans by inserting earplugs and removing them after a period (typically days or weeks). Like cochlear synaptopathy/HHL, CHL has been associated with a homeostatic increase in central gain in the auditory pathways (Sheppard et al., 2018; Hutchison et al., 2023), and a concomitant increase in the subjective experience of tinnitus (Zeng, 2013). Compared to permanent lesions of the sensory surface of the inner ear, HHL and CHL provide a means of exploring the extent to which principles of coding efficiency can explain changes, including potentially counterintuitive changes, in neural coding capacity and listening performance as a consequence of altered sensory input.

Here, we test whether principles of coding efficiency can account for altered sensory representations of unfolding sound-level distributions following noise exposure designed to induce HHL and ear-plugging designed to generate CHL. Specifically, we assess whether a recent theoretical framework (Młynarski et al., 2021) that combines the normative theory of optimal coding (sensory systems evolving to solve essential tasks and processes) with statistical inference that ignores *a priori* notions of biological function, can explain the level of optimality achieved by auditory neurons when rapidly adapting to unfolding distributions of sound intensities, including when coding capacity is altered by HHL or CHL. Employing this framework allows us to differentiate and contrast these two seemingly unrelated pathologies using common informational units, supporting an understanding of sensory deficits in terms of changes in the dynamics by which information is encoded in the auditory brain. Employing previously published (Monaghan et al., 2020) and unpublished *in vivo* recordings, we show that up-regulation of neural gain (the consequence of loss of selective afferent inputs in synaptopathy/HHL or reduced sensory input gain in CHL) alters the context-dependent modulation of auditory-midbrain coding, with the clearest effects appearing in quiet acoustic environments where the information-cost trade-off most clearly differentiates lesion conditions. This framework helps reconcile the potentially counterintuitive observation *in vivo* that whilst neural coding of speech sounds in background noise and adaptation to the statistics of sound-intensity distributions are impaired in HHL compared to controls at relatively high intensities, they can be better than normal at quiet-to-moderate intensities. This framework can also account for changes in neural coding capacity that arise from rapid changes in input gain; here, specifically following removal of earplugs designed to induce CHL.

Together, our data suggest that an optimal-coding framework developed to explain sensory representations is useful for exploring and explaining communication problems currently not explained in terms of standard clinical manifestations such as permanently elevated hearing thresholds. Applying fundamental principles of computational neuroscience to subtle sensory deficits offers a phenomenological level of abstraction to investigate normal and impaired sensory systems in a continuum of optimality.

## Results

We examined neural responses recorded from populations of neurons recorded in the inferior colliculus to acoustic contexts defined by their statistical distribution of sound intensities. Specifically, we assessed adapted neural thresholds (i.e., the lowest intensities from a defined distribution that evoked a significant increase in firing rates above the midpoint between its minimum and maximum) and gain (slope) parameters from rate-vs.-intensity functions (RIFs). We then constructed context-specific optimization priors over threshold-slope parameters using a utility function that trades mutual information against a spiking-cost term, following the method of Młynarski et al. (2021). Finally, we infer for each group and context the hyperparameters that best explain the observed distributions of thresholds and gains, yielding trajectories in a common information-cost coordinate system that quantify how each lesion disrupts efficient coding across acoustic contexts (HPRs). This allowed us to compare directly the effects of two subtle pathological states, HHL and CHL, on the dimensions of coding utility, entropy, and central neural gain. The data are presented in the context of the two pathologies, each with two groups of animals: sham-exposed (SH; controls) and noise-exposed (NE) for HHL, and ear-plugged (EP) and post-plug removal (PP) for CHL. The HHL dataset employed here corresponds to additional experiments in the same recording sessions of Monaghan et al. (2020), whereas the CHL dataset has remained unpublished to date.

### Neural adaptation to acoustic contexts as a substrate for solving the dynamic range problem

Neural responses were recorded to statistically defined distributions of sound intensities (acoustic contexts), each characterized by a high-probability region (HPR; see Figure 1 A, D) spanning 12 dB SPL, from which 80% of stimulus presentations were selected, with the remaining 20% selected from other intensities in the range 24-96 dB SPL. A uniform distribution (no HPR) was also presented. The stimulus consists of concatenated tokens of white noise that change in intensity every 50 ms, the specific sequence representing a realization of the acoustic context (the HPR distribution, Figure 1 A). These stimuli have been used to explore neural mechanisms contributing to dynamic-range coding in the auditory nerve (Wen et al., 2009), midbrain (Dean et al., 2005, 2008; Bakay et al., 2018), and cortex (Watkins and Barbour, 2008, 2011). Neural responses to them are modulated by the mean intensity of the stimulus distribution (Auerbach and Gritton, 2022), with the operating range of the RIFs shifting with the underlying HPR distributions (Dean et al., 2005, 2008). This capacity for RIFs to adapt to the underlying distribution of sound intensities is suggested to contribute to the exquisite (1 dB) discrimination thresholds that hold over 12 orders of magnitude (120 dB, (Viemeister, 1988; Nizami and Schneider, 1997; Auerbach and Gritton, 2022)).

**Figure 1.**
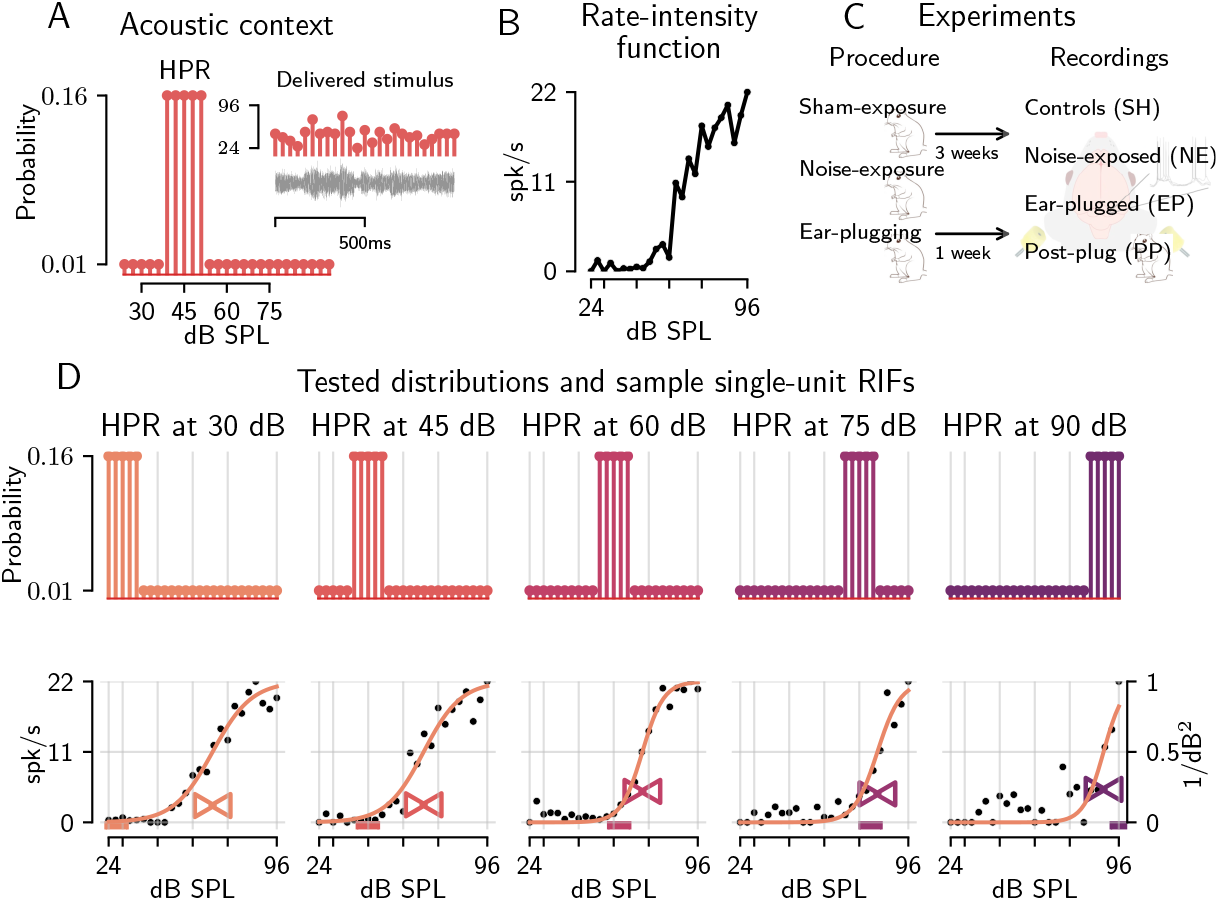
**(A-B)** Acoustic context and example single-unit rate-intensity functions (RIFs). In A, horizontal coloured bars mark the 12-dB HPR window for each context. In B, one example RIF is shown for the stimulus in A. **(C)** Depiction of the experiments’ timeline for the two pathological conditions (HHL and CHL). HHL condition consists of sham-exposed (SH; controls) and noise-exposed (NE), CHL condition of ear-plugged animals recorded pre-(EP) and post-plug (PP) removal. All recordings were made in the IC, three weeks after the procedures of noise-exposure and one week after ear-plugging. **(D)** Shifting of RIF center in a sample single-unit for each acoustic context with a sigmoidal fit. Bow-ties mark the fitted threshold (x-coordinate) and gain (y-coordinate), read from the auxiliary axes at the right of panel D.

RIFs evoked by HPR stimuli differ in adapted threshold and gain along the auditory pathway, with auditory-nerve responses showing lower gain (i.e., less steep) thatn the progressively steeper functions observed in midbrain and cortex (Wen et al., 2009; Dean et al., 2005, 2008; Watkins and Barbour, 2008, 2011; Auerbach and Gritton, 2022). Sensorineural hearing loss triggers two main effects: RIF thresholds shift and central neural gain in IC neurons is up-regulated, increasing the steepness of midbrain and auditory cortex RIFs, and elevating the firing-rate in response to louder sounds (Syka et al., 1994; Auerbach and Gritton, 2022). The IC is the major auditory nucleus of the midbrain and a virtually obligatory synaptic station in the ascending and descending auditory pathways. Considering this, the IC is a natural locus in which to assess adaptation to complex sounds (Nelken, 2014; Chechik et al., 2006; Peng et al., 2024).

IC neural responses to six different acoustic contexts, five HPR stimuli (30, 45, 60, 75, and 90 dB SPL) and a uniform distribution of sound intensities spanning the full 24-96 dB range (Figure 1 D), were recorded in animals subjected to the two pathological conditions HHL and CHL. In the HHL condition, noise-exposed (NE) animals were anaesthetized and exposed to an octave-band noise (2-4 kHz, 105 dB SPL) whose center frequency, bandwidth, intensity, and duration were previously determined to elicit synaptopathy/HHL (Kujawa and Liberman, 2009; Bakay et al., 2018); whilst the sham-exposed (SH) group were subject to anesthesia only (Figure 1 C). IC neural recordings were conducted after NE animals recovered their temporary elevated thresholds to normal thresholds (approximately three weeks following noise exposure). In the CHL condition, animals were anaesthetized and subject to the insertion of earplugs bilaterally, with IC neural recordings made one week later before the earplug was removed (EP) and immediately post removal (PP), Figure 1 C.

Evoked responses to the six different acoustic contexts (five HPRs and a uniform distribution; Figure 2 and Figure 3 A) indicate the dependence of the average adapted thresholds on the centers of the HPRs (mean sound intensity). Fisher information (FI, Figure 2 and Figure 3 A&B) provides a measure of neural coding accuracy and reflects the steepness of the RIFs, counterweighted by variance of the firing rate. It has been used to quantify the sensitivity of neural responses to small changes in stimulus intensity, with higher FI indicating greater encoding precision (Dean et al., 2005; Bakay et al., 2018). Here, we use FI as a descriptive metric of sensitivity to facilitate comparisons with published reports of dynamic-range adaptation, but we rely on the framework of optimization priors (outlined below) to quantify system-level trade-offs between information and metabolic costs inaccessible to FI analysis. RIFs were then normalized and fitted to a sigmoid with two parameters (threshold and gain), reducing the dataset to a two-dimensional data set (bottom of Figure 2 and Figure 3 B). The distribution of values in the data took different forms depending on the HPR, and these differences can be observed by exploring the marginals of the threshold-vs.-gain distributions (above and at the right of each scatter plots).

**Figure 2.**
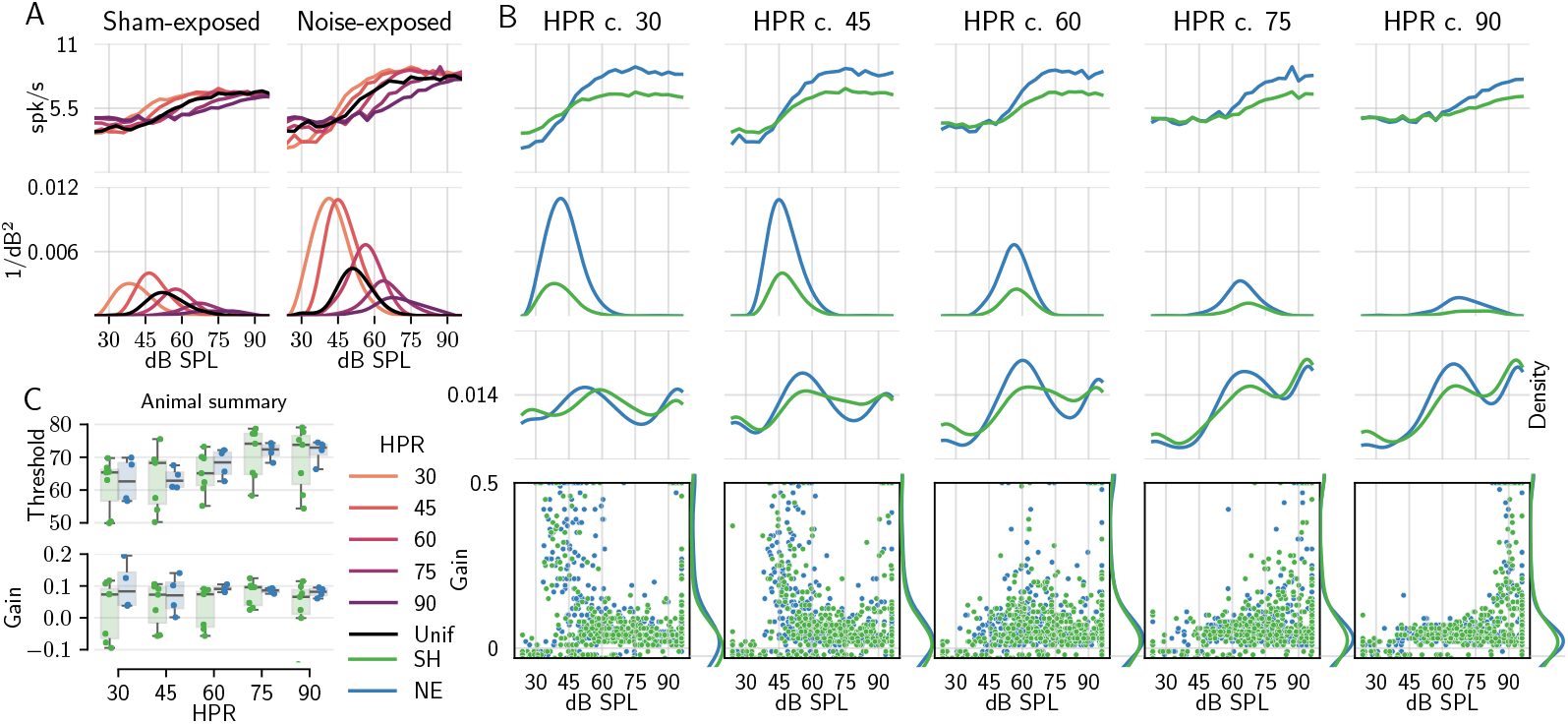
Mean-sound level adaptation in the HHL condition **(A)** Average population-response RIFs (first row) and Fisher information (second row), organized by groups (columns). **(B)** Average population responses organized by HPR (columns). RIFs and Fisher information are as in A (first two rows), while the bottom row shows thresholds and gains of individual neurons, with density at top and left. **(C)** Animal-level summary of mean adapted thresholds and gains across HPRs for sham-exposed (SH) and noise-exposed (NE) animals.

**Figure 3.**
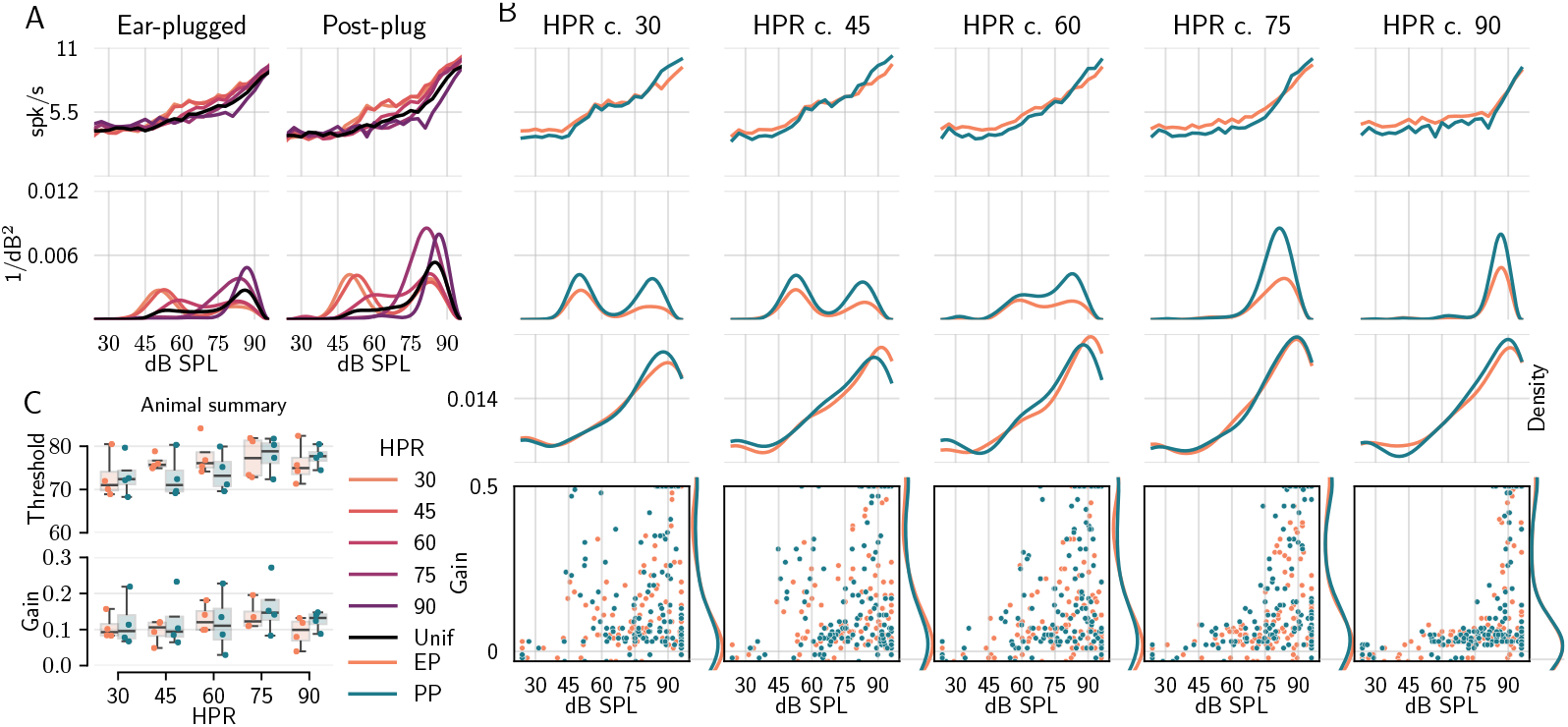
Mean sound level adaptation in the CHL condition. **(A)** Average population-response RIFs (first row) and Fisher information (second row), organized by group (columns). **(B)** Average population responses organized by HPR (columns); the bottom row shows thresholds and gains of individual neurons, with density at top and left. **(C)** Animal-level summary of mean adapted thresholds and gains across HPRs for ear-plugged (EP) and post-plug (PP) animals.

#### Hidden hearing loss alters the adaptive response of midbrain neurons to unfolding sound environments

We first explored the impact of HHL on the neural representation of sound environments characterized by their statistical distribution of sound intensities. Auditory nerve fibers implicated in HHL (the high-threshold, low spontaneous-rate fibers) are thought to modulate sensory gain through a series of reflexive brainstem pathways (Kujawa and Liberman, 2009; Bharadwaj et al., 2014; Johannesen et al., 2022). Damage to these fibers spares hearing thresholds but leads to an overall reduction in neural activity at suprathreshold sound intensities, potentially altering neural gain in downstream auditory neurons. Consistent with this interpretation, responses of IC neurons in noise-exposed (NE) animals showed evidence of steeper fitted RIFs for quiet to moderately quiet acoustic contexts (i.e., HPRs 30 dB SPL and 45 dB SPL), with lower firing rates at low intensities and greater firing rates at higher, relative to sham-exposed (SH) animals (Figure 2 B). This difference in firing rates between NE and SH animals was reduced as the general acoustic context became louder (HPRs 60 dB SPL and 75 dB SPL). Quantified in terms of FI, intensity coding was more pronounced in NE compared to SH animals for overall quieter environments (HPR 30 dB in Figure 2 B) but was progressively less distinguishable in moderate to loud contexts (HPRs 45 dB, 60 dB and 75 dB).

At the animal level (Figure 2 C), adapted thresholds in NE tended to be lower at the quietest HPR, but this effect was modest and the group × HPR interaction for thresholds was not significant (Hedges’ *g* = 0.29, 95% CI [− 0.84, 1.42], *p* = 0.72 at HPR 30; threshold modulation across contexts: *p* = 0.920, FDR corrected; Table 1). The gain summary in Figure 2 C was more distinctive: NE and SH separated increasingly at louder HPRs, with sham-exposed animals showing progressively greater gain increases than noise-exposed animals. A linear mixed-effects model confirmed that gain modulation across acoustic contexts differed significantly between NE and SH groups (group × HPR interaction: *χ*^2^(4) = 16.66, *p* = 0.009, FDR corrected; Table 1), with sham-exposed animals showing progressively greater gain increases relative to noise-exposed animals at louder HPRs. This context-dependent convergence of NE gain distributions was additionally supported by a significant group × HPR interaction on distributional spread (absolute gain deviation from the group-and-HPR median: *χ*^2^(4) = 10.03, *p* = 0.040, FDR corrected; Table 1), consistent with NE neurons converging on similar elevated gain states at louder contexts where surviving low-threshold fibers dominate. In contrast, an informative null result was obtained for threshold modulation: the group × HPR interaction for adapted thresholds was not significant (*p* = 0.920, FDR corrected), indicating that the HHL empirical effect is localized in gain rather than in threshold adaptation.

**Table 1.** Table S1. Empirical group comparisons (linear mixed-effects models). Rows sorted by *p*FDR. Interaction models report joint Wald *χ*^2^ (4); main-effect models report coefficient *β*. ** = confirmed effect after Benjamini-Hochberg false-discovery-rate correction (*p*_FDR_ *<* 0.05); * = marginal trend (0.05 ≤ *p*_FDR_ *<* 0.10).

| ID | Claim (Groups) | Model | $p_{raw}$ ($p_{FDR}$) |
| --- | --- | --- | --- |
| E8 | CHL thresholds exceed controls (EP vs SH) | threshold ~ group × HPR + (1\|animal);<br>$\chi^2(4) = 17.94$ | 0.001 (0.004)** |
| E9 | CHL thresholds exceed synaptopathy (EP vs NE) | threshold ~ group × HPR + (1\|animal);<br>$\chi^2(4) = 15.43$ | 0.004 (0.006)** |
| E1 | Gain modulation differs across contexts (NE vs SH) | gain ~ group × HPR + (1\|animal);<br>$\chi^2(4) = 16.66$ | 0.002 (0.009)** |
| E_SP1 | Gain distributions converge at loud (NE vs SH) | \|gain - median\| ~ group × HPR +<br>(1\|animal); $\chi^2(4) = 10.03$ | 0.040 (0.040)** |
| E_SR1 | Spontaneous rate lower at quiet (NE vs SH) | spont_rate ~ group × HPR + (1\|animal);<br>$\chi^2(4) = 11.26$ | 0.024 (0.048)** |
| E10 | Post-plug thresholds above controls (PP vs SH) | threshold ~ group × HPR + (1\|animal);<br>$\chi^2(4) = 9.42$ | 0.051 (0.051)* |
| E_SP2 | Threshold distributions narrow at loud (NE vs SH) | \|thresh - median\| ~ group × HPR +<br>(1\|animal); $\chi^2(4) = 10.66$ | 0.031 (0.061)* |
| E_SR2 | EP spontaneous rate elevated (EP vs PP) | spont_rate ~ condition + (1\|animal);<br>$\beta = -0.47$ | 0.093 (0.464) |
| E_DR1 | Dynamic range wider across contexts (NE vs SH) | rate_range ~ group × HPR + (1\|animal);<br>$\chi^2(4) = 3.05$ | 0.549 (0.824) |
| E_FI2 | Localized FI higher at HPR 30 (NE vs SH) | log(FI<sub>24-50</sub>) ~ group + (1\|animal);<br>$\beta = -0.37$ | 0.277 (0.831) |
| E6 | Ear plug elevates spike rate (EP vs PP) | rate ~ condition + (1\|animal); $\beta = -0.18$ | 0.542 (0.904) |
| E4 | Threshold modulation differs across contexts (NE vs SH) | threshold ~ group × HPR + (1\|animal);<br>$\chi^2(4) = 2.25$ | 0.690 (0.920) |
| E_FI1 | Fisher information higher at quiet (NE vs SH) | log(total_FI) ~ group × HPR + (1\|animal);<br>$\chi^2(4) = 0.64$ | 0.959 (0.959) |
| E_FI3 | PP coding precision recovers uniformly (EP vs PP) | log(total_FI) ~ condition × HPR +<br>(1\|animal); $\chi^2(4) = 0.34$ | 0.987 (0.987) |
| E5 | Ear plug elevates thresholds (EP vs PP) | threshold ~ condition + (1\|animal);<br>$\beta = -0.83$ | 0.431 (1.000) |
| E7 | Plug removal alters gain modulation (EP vs PP) | gain ~ condition × HPR + (1\|animal);<br>$\chi^2(4) = 0.67$ | 0.955 (1.000) |

#### Conductive hearing loss reshapes adaptive coding and enhances gain control in midbrain neurons

Identical analyses to those applied to neural recordings from animals with HHL were undertaken for animals subject to conductive hearing loss (CHL). CHL is characterized by a reduction in input gain to the inner ear. Here, CHL was induced by transient ear-canal plugging, which we treat as a reversible attenuation of sound level reaching the inner ear. This manipulation provides a simple way to compare coding before and immediately after restoration of input level.

Neural recordings were made with the ear plug (EP) in place and immediately following its removal (PP). For all HPRs, neural firing rates were overall higher in the EP compared to the PP condition (Figure 3 B; pooled-neuron: *p* < 0.001; animal-level: Hedges’ *g* = 0.30, 95% CI [− 0.92, 1.51], one-tailed directional *p* = 0.33, *N* = 4 vs 4 animals; linear mixed-effects model: *p* = 0.904, FDR corrected; direction-consistent but not confirmed; Table 1), consistent with a reduction in input gain generating an over-compensatory increase in internal, neural gain. Evident also in Figure 3 B (second row) is the bimodal nature of the distribution of FI in neural recordings from both EP and PP conditions. The bimodal FI profile likely reflects two distinguishable subpopulations visible in the threshold-gain distributions: a low-threshold, high-gain group that concentrates coding accuracy near conversational levels, and a high-threshold subgroup (consistent with the distinct population described by Dean et al. (2005)) that provides a secondary peak at louder intensities. This high-threshold peak is absent for the uniform distribution, in which probability mass is spread evenly and there is less statistical incentive to maintain a specialized loud-coding population.

Specifically, coding accuracy peaks for intensities in the range 45-60 dB SPL and in the range 75-90 dB SPL for lower HPRs (30–60), while for HPR 75 and HPR 90 only a single peak at high intensities was present. FI was low in the 60–75 dB SPL range for HPR 30 and HPR 45, where the two peaks were separated by a region of diminished coding; for HPR 60 this valley was absent and FI peaked in this range instead. Bimodality was more pronounced for PP than for EP for most HPRs, with higher FI (greater coding accuracy) at higher intensities. For instance, for HPR 30, encompassing 24-36 dB SPL, an intensity of ∼80 dB SPL has a low probability of appearing but, nevertheless, FI peaks near this value. These two peaks in FI remained largely fixed around the same intensities (50 and 80 dB SPL) independent of the HPR, with the lower peak shifting upward to ∼60 dB SPL for HPR 60 and disappearing for HPR 75 and HPR 90. Interestingly, similar peak firing rates were generated for all stimulus conditions in both EP and PP states, but notably the spontaneous firing rate (firing rates at sound levels below adapted thresholds for each HPR and the uniform distribution) were higher in the EP compared to the PP state. This pattern is consistent with a shift in gain state after plug removal, but with only partial normalization over the timescale of the recording session.

The animal summary in Figure 3 C shows that thresholds remained elevated across HPRs in both EP and PP, whereas gain differences between conditions were smaller and more variable across animals. Unlike in HHL, conductive attenuation produced by the ear plug reduces the effective input level across frequencies, and this reduction cannot be fully offset by a central gain increase over the timescale of the recording session. Consequently, for both the EP and PP conditions, adapted thresholds were substantially higher than in HHL conditions. Cross-cohort linear mixed-effects models confirmed that EP thresholds exceeded both sham controls (group × HPR interaction: *χ*^2^(4) = 17.94, *p* = 0.004, FDR corrected) and noise-exposed animals (*χ*^2^(4) = 15.43, *p* = 0.006, FDR corrected), with this elevation largest in quiet-to-moderate contexts and narrowing at louder HPRs (Table 1), indicating that conductive attenuation displaced the effective operating range of IC responses. Post-plug thresholds remained above sham controls but the group × HPR interaction was marginal (group × HPR interaction: *p* = 0.051, FDR corrected), indicating incomplete but directionally consistent threshold renormalization over the timescale of the recording session.

### Optimization priors for acoustic contexts

In clinical practice, hearing is assessed by measuring behavioural sensitivity (thresholds) to pure tones presented at a range of frequencies, with the lowest audible intensity generating the audiogram. This gold-standard assessment of hearing function is inherently insensitive to supra-threshold encoding quality and cannot distinguish how well a subject can listen in noisy environments or selectively attend to a particular sound source (Plack et al., 2014; Bharadwaj et al., 2014; Liu et al., 2024a,b). Supra-threshold listening difficulties are commonly framed as limitations in processing temporal envelope and temporal fine-structure cues, and several methodologies have been used to measure these in clinic, such as speech-in-noise recognition, amplitude-modulation detection, temporal gap detection, frequency/pitch discrimination, and binaural temporal fine-structure tests (e.g., interaural phase/time sensitivity) (Moore, 2008; Hopkins and Moore, 2011; Bernstein et al., 2013; Kortlang et al., 2016; Zaar et al., 2023, 2024; Carney, 2018; Johannesen et al., 2016). However, these measures show significant inter-subject variability in performance and are rarely employed in clinical practice (Carcagno and Plack, 2022; Sanchez-Lopez et al., 2018, 2020; Lutfi et al., 2020, 2021). Moreover, despite substantial evidence from animal and human work that cochlear synaptopathy and related deafferentation can degrade supra-threshold coding while leaving audiometric thresholds largely normal, the lack of sensitive, reliable, and standardized clinical biomarkers has made the assessment and diagnosis of “hidden hearing loss” in everyday clinical settings particularly challenging (Kujawa and Liberman, 2009; Plack et al., 2014; Bramhall et al., 2019; Valderrama et al., 2018; Johannesen et al., 2019; DiNino et al., 2022; Valderrama et al., 2022; Liu et al., 2024a,b).

Here we take an information-theoretic approach that explicitly quantifies how much information about complex, noisy acoustic environments is conveyed by neural responses, and how that information is traded off against the metabolic costs of spiking. Młynarski et al. (2021) described a framework that combines normative brain theories with statistical inference in a Bayesian manner. Here, we use this framework to test hypotheses concerning the degree of optimality achieved by IC neurons in animals subject to HHL or CHL. Briefly, the framework first evaluates a utility function *U* (*θ*) on simulated responses of a range of model neurons (with two parameters *θ* = {*x*_0_, *k*}, threshold and gain), then transforms that utility surface into a probability distribution *P* (*U* (*θ*)), written as *P* (*θ*) for simplicity. This is then used as a Bayesian prior to which the likelihood of a dataset 𝒟 (in this case, the adapted thresholds and gains of RIFs recorded from IC neurons) can be added to calculate a log-posterior (hence the name optimization prior, OP), i.e.,

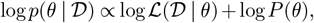

When optimization-prior quantities are compared across groups or contexts below, we use the hierarchical animal-bootstrap procedure described in Section rather than pooled-neuron significance tests. The utility function is defined as a trade-off between information transmission and metabolic costs, where the former is quantified by the mutual information *I* between the stimulus category vector *c*_*t*_ and the response vector *r*_*t*_, and the latter is represented as a penalty proportional to the average firing rate of the neuron:

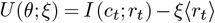

*ξ* is a hyperparameter that weights the firing-rate penalty and therefore determines whether utility is dominated more by information transmission or by metabolic cost. Throughout, we compute mutual information using a binary response variable *r*_*t*_ ∈ {0, 1} obtained by thresholding the normalized response (equivalently, a spike/no-spike indicator per 50-ms bin). In this representation, ⟨*r*_*t*_⟩ represents a spike probability per bin and serves as a convenient proxy for metabolic cost following normalization. This function is then transformed into a maximum entropy distribution, serving as a prior in a Bayesian framework:

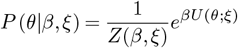

where *Z*(*β, ξ*) is a normalization constant. This form is the unique maximum-entropy distribution subject to a constraint on expected utility (Jaynes, 2003), ensuring the prior encodes the normative utility surface without imposing additional structural assumptions. With this convention, larger *β* concentrates prior mass on higher-utility parameter settings, whereas *β* = 0 yields a uniform distribution over the discretized parameter grid. The family of OPs *P* (*θ* | *β, ξ*) are Boltzmann distributions that maximize the entropy among all distributions with the same average utility, defined as:

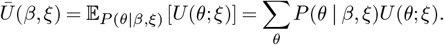

*β* is an *optimization hyperparameter* that controls how structured the OPs are: *β* = 0 renders a uniform distribution, indicating the same utility for all parameters *θ* in the grid (stimulus independent), and as *β* → ∞, it yields OPs that are increasingly condensed around an optimal region (stimulus dependent).

To yield OPs for each HPR, we followed the procedure of Młynarski et al. (2021), but using the HPRs as stimuli (Figure 4). Although we provide a mathematical description for a single HPR distribution, the same derivation applies to all five (centered at 30, 45, 60, 75, and 90 dB SPL), where *p*_*s*_(*x*) is defined as the HPR-distribution for intensity *x* (dB SPL). We categorize each intensity sample at each time point into one of three regions below (b), in (i) and above (a), the HPR (i.e., bHPR, iHPR, and aHPR), generating a category vector *c*_*t*_ from *s*_*t*_, (Figure 4 A). For each HPR, we generated realizations of 40,000 samples, resulting in five separate stimulus vectors *s*_*t*_, subsequently quantized into *c*_*t*_, (Figure 4 B). Model neurons were determined by parameters *θ* = {*x*_0_, *k*}, where *x*_0_ is the threshold and *k* is the gain, quantized in a grid of 25 × 25 values for *x*_0_ and *k* respectively, Figure 4 C. Inputting *s*_*t*_ into the logistic nonlinearity 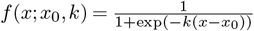 yields a response vector *r*_*t*_(*θ*) for all *θ* = {*x*_0_, *k*}. From these simulated response vectors, we computed the sample mean *µ*_*r*_ = ⟨*r*_*t*_⟩ and the mutual information *I* (*c*_*t*_; *r*_*t*_), defined as

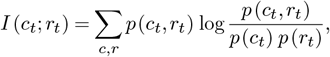

where the joint and marginal distributions were estimated from 2D histograms with three stimulus categories (bHPR, iHPR, aHPR) and two response states ({0, 1}, threshold at 0.5). Because *r*_*t*_ is binary, the mutual information here quantifies how well spike/no-spike responses discriminate the three stimulus categories, and the cost term ⟨*r*_*t*_⟩ corresponds to spike probability per bin. The binary response with three stimulus categories follows the worked example of Młynarski et al. (2021), where this representation was adopted as the most parsimonious encoding that preserves the category-discrimination structure of the utility; finer response or stimulus discretizations would increase MI monotonically but without changing the qualitative geometry of the optimization-prior landscape. For each HPR distribution, the mutual information (Figure 4 E, top row) is maximal near the upper and lower boundaries of the HPR, indicating the most informative regions for decoding the stimulus into the three categories. Conversely, the mean firing rate *µ*_*r*_ (Figure 4E, bottom row), peaks below the HPR region for neurons with positive gain. Neurons with negative gain display the opposite pattern: elevated baseline firing that declines with increasing intensity.

**Figure 4.**
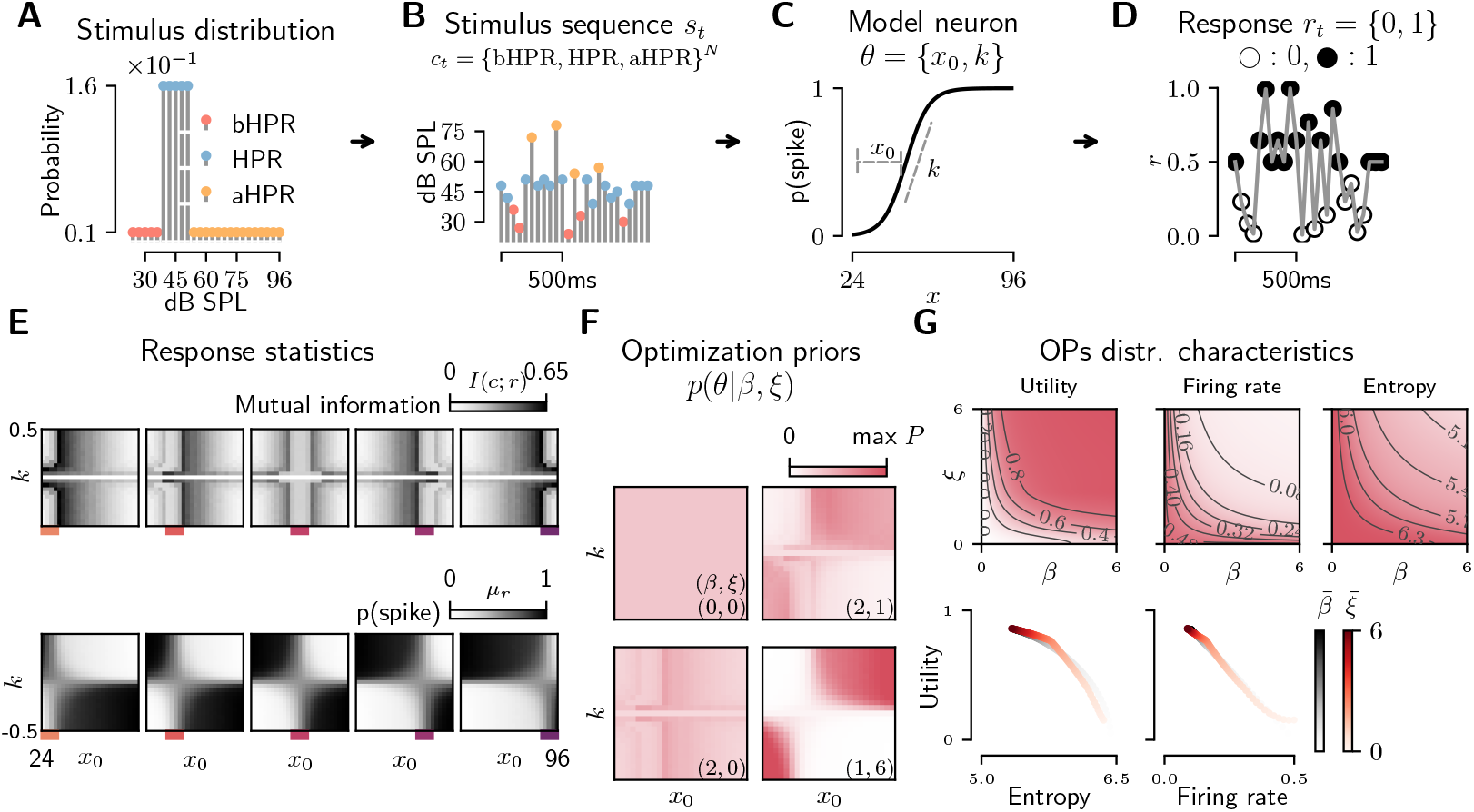
**(A-D)** Simulation of responses of model neurons. First, a stimulus sequence is generated (B) from the distribution in (A), later inputting the sequence into a model neuron (C) with certain threshold (*x*_0_) and gain (*k*); the output of the model neuron is binarized to construct a binary spike train (D), used to calculate statistics associated with its parameters (*θ*). The same process is repeated for a grid of model neurons. **(E)** Statistics (for each HPR) of the grid of model neurons, mutual information (upper panel) and average spike rate (bottom panel). These two statistics are the elements by which the utility function is constructed. **(F)** Four examples of OPs *P* (*θ* | *β, ξ*), visualized for different combinations of (*β, ξ*), illustrating how utility trade-offs modulate the structure of the priors; (0, 0) renders a uniform distribution, for moderate *β* = 2 and increasing *ξ*, the OPs take the shape of the inverse of the firing-rate rather than being driven by the mutual information (lower values of *ξ*). **(G)** Distributional characteristics of OPs as a function of (*β, ξ*). The upper panel shows three contour plots that summarize the effect of hyperparameters on the OPs. Here utility refers to the normalized average utility 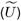, entropy to *H* [*P* (*θ* | *β, ξ*)], and firing rate to *µ*_*r*_, averaged across the grid of neurons. The iso-contour lines shows different combinations of (*β, ξ*) that yield the same utility, entropy, or firing rate, highlighting their exponential shape. The bottom panel plots the relationship between utility vs entropy and firing-rate, averaging over *β* and *ξ*.

Hyperparameters *β, ξ* determine one OP inside the family of OPs, controlling their core statistics such as average utility, entropy, and mean firing-rate. In general, the average utility grows monotonically with *β* and reaches maximum when *β* → ∞. We were particularly interested in comparing OPs within a family of them, so we define a normalized average utility as

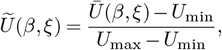

where *U*_min_ and *U*_max_ represent the lowest and highest attainable average utility values, respectively (among all *β* and *ξ*).

We additionally computed the entropy and average spike rate of each OP *P* (*θ* | *β, ξ*) across the grid of hyperparameters to evaluate the diversity and metabolic demand of the induced priors as:

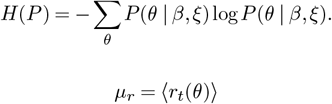

Both entropy and mean firing rate decrease as *β* increases. When *β* = 0, the prior *P* (*θ* | *β, ξ*) is uniform, resulting in maximum entropy. Similarly, the mean firing rate decreases with increasing *β*, reflecting a narrower distribution and reduced variability in the parameter space.

### Inference of optimization priors and lesion-specific neural coding trajectories across acoustic contexts

Taking the sets 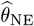 and 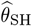 of fitted neural parameters (pairs of thresholds and gains) as instances (samples) of systems that operate under the normative model into consideration, we then estimate the hyper-parameters of the optimization prior that are most likely to have generated the data (threshold-gain distributions). This is achieved by hierarchical maximum *a posteriori* (MAP) estimation, where the posterior 𝒫 over the two hyper-parameters was approximated as

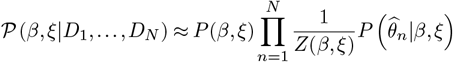

where {𝒟_1_, …, 𝒟_*N*_} is the set of each neuron spike-train data 𝒟_*n*_. This expression makes use of both simplifications described in section 4.2, namely *data-rich* and *multiple instances simplification*. The data-rich simplification highlights that, in this case, there is sufficient data from each neuron such that the likelihood distribution of neural parameters is narrowly distributed and that, therefore, the procedure can be reduced to function fitting (sigmoid). Multiple instances of the system are also available in the form of multiple neurons simultaneously recorded, which constructs the datasets for each animal group as threshold-gain samples.

MAP estimation gives a point estimate of *β* and *ξ* (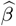 and 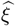, respectively), by choosing the hyper-parameters that maximise the marginal posteriors, i.e.,

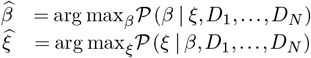

The vertical maxima in Figure 5 A reflect these MAP estimates, for all HPRs and both animal groups, using data from all neurons.

**Figure 5.**
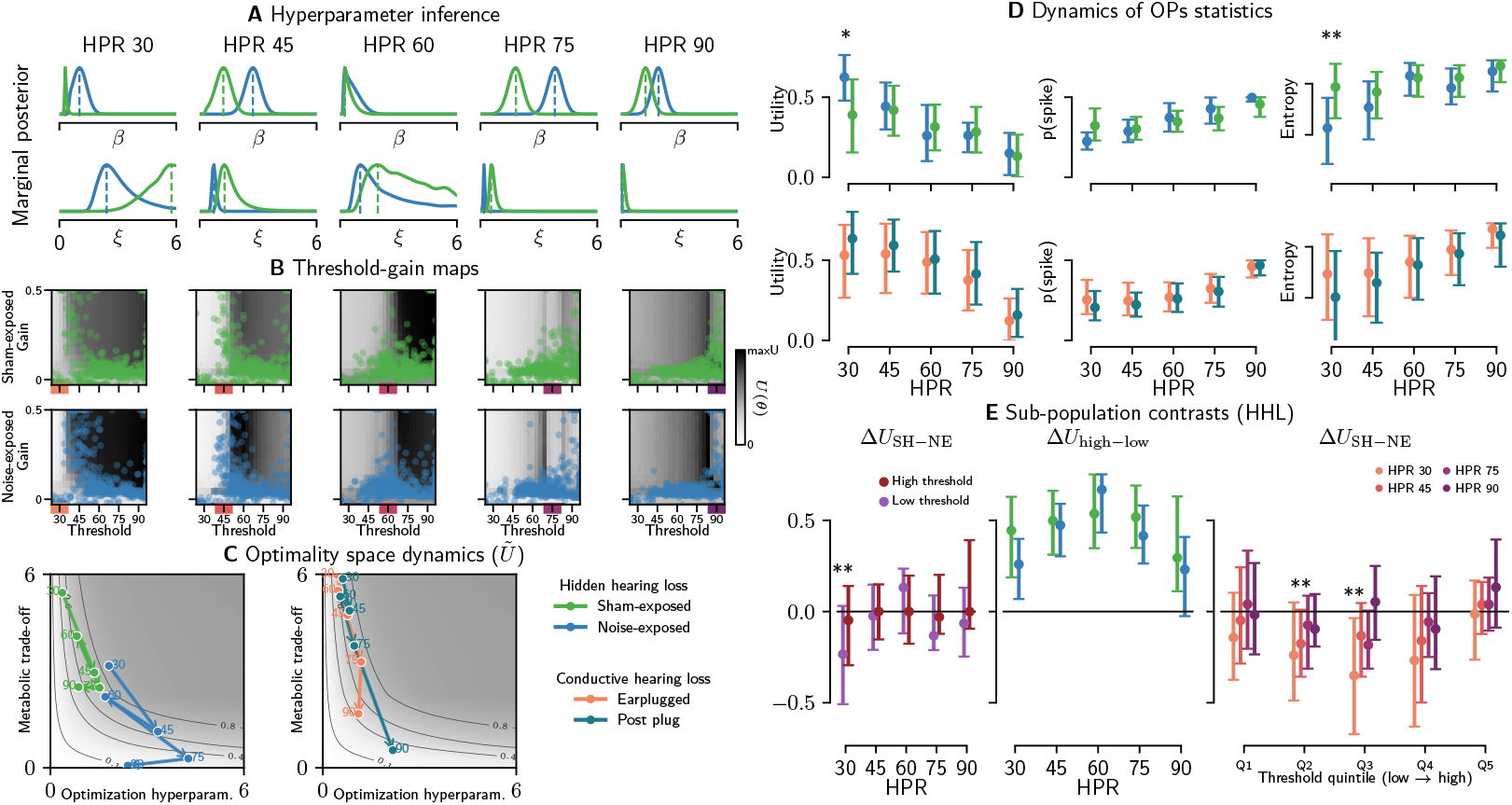
Population hyperparameter inference. **(A-B)** Example of maximum a-posteriori estimation. For each acoustic context, the posterior distribution was calculated (plotted in panel A) from the set of thresholds and gains, and the values of *β* and *ξ* with the maximum posterior were selected. Each pair (*β, ξ*) determines an optimization prior that is associated with the dataset, which is plotted in the background of panel B, along with the threshold-gain datasets for each group (HHL condition). Overlaying the datasets on the corresponding priors illustrates how closely the empirical distributions follow the predictions of the optimization framework. **(C)** Dynamics of the hyperparameter space. Arrows show how inferred optimization hyperparameters move across acoustic contexts (HPRs), separately for HHL and CHL, over a normalized-utility background. Arrow labels mark HPR centers. **(D)** Statistics of the associated optimization priors for each animal group. Each point shows the mean obtained across hierarchical animal-bootstrap replicates using the same inference procedure; error bars indicate 95% confidence intervals. Utility, spike probability, and entropy are plotted on their own axes. **(E)** Two-population contrasts for HHL (split at 85 dB SPL median adapted threshold). Left, SH-NE utility contrast by HPR for low- and high-threshold populations. Middle, within-group high-minus-low utility differences by HPR. Right, threshold-resolved Δ*U*_SH−NE_ across across-context threshold quintiles, one series per HPR; markers denote Pr(NE *>* SH) ≥ 0.95 (**) and ≥ 0.99 (***).

To quantify uncertainty while reducing unit-level pseudo-replication, we repeated the inference using hierarchical bootstrap resampling at the animal level (45 threshold-gain samples per HHL replicate and 30 per CHL replicate; averages are shown in Figure 5 B-E). For visualization, Figure 5 B plots the same distributions of threshold-vs.-gain as in Figure 2 B, but now with the matched optimization prior distribution represented by the shaded background. From this, it is observed that the samples of thresholds and gains tend to cluster near the fringes of the HPR, although the resulting group contrasts are strongest in quiet contexts and become progressively less stable as mean intensity increases.

Across acoustic contexts, the inferred hyperparameters (*β, ξ*) trace trajectories in the optimization-prior space, visualized in Figure 5 C as arrows progressing from quieter to louder HPRs (30→45, 45→60, 60→75, and 75→90 dB SPL). Here, *β* controls the concentration or organizational structure of the prior over threshold-gain parameters (lower entropy and tighter mass near high-utility solutions as *β* increases), whereas *ξ* sets the strength of the metabolic penalty on the mean firing rate (larger *ξ* enforces lower rates). Panel C shows these trajectories over normalized-utility contours, whereas Figure 5 D summarizes the corresponding utility, spike probability, and entropy on separate axes.

For HHL groups, the first arrow segment in Figure 5 C (30→45 dB SPL) points down and to the right, indicating increases in *β* and reductions in *ξ*; CHL trajectories showed only marginal changes over this transition. This implies that the system relaxes the firing-rate constraint while organizing the prior more tightly around informative threshold-gain configurations. The subsequent 45→60 dB SPL transition reverses this pattern, with lower *β* and higher *ξ*, and these segments still lie near iso-contours, consistent with coordinated trade-offs in a transitional regime. At 60→75 dB SPL, *ξ* decreases again in all groups and trajectories generally show increases in *β* (rightward), signaling renewed movement toward a lower-constraint regime with higher firing rates and a concomitant drop in normalized utility. At 75→90 dB SPL, *ξ* falls further and normalized utility declines again, while implied firing rises sharply; for HHL, especially NE, the loudest context therefore lies close to a boundary-dominated low-*ξ* regime. Relative positioning of trajectories reveals pathology-specific operating regimes across contexts.

Across 30 to 75 dB SPL, responses from animals subjected to CHL (ear-plugged, EP and post-plug, PP) occupy lower *β* and higher *ξ* than noise-exposed animals at matched HPRs, with PP showing slightly higher utility than EP and, in several contexts, slightly higher *β*. CHL trajectories remain relatively localized along iso-contours up to HPR 75, reflecting a compressed dynamic range; at the 75→90 transition, particularly for PP, trajectories shifted sharply toward lower ξ in a manner comparable to HHL. Within HHL, controls (SH group) tend to remain at higher *ξ* than NE across contexts, whereas differences in *β* are more context dependent. Trajectories in NE animals begin at lower *ξ* and higher *β* than those of SH at 30 dB and undergo a further reduction in *ξ* with a rise in *β* over 30→45 dB, keeping NE in a lower-penalty regime than sham at the same HPR. Across 45→60 dB both NE and SH groups show decreases in *β*; *ξ* increased substantially for SH but only marginally for NE, and this transition should be interpreted as a context-dependent trend rather than a fixed reversal. By 75 dB, both groups have returned toward a lower-*ξ* regime, and by 90 dB utility differences are small while NE lies closer to the *ξ* = 0 boundary.

These movements in (*β, ξ*) space systematically modulate the statistics of the matched optimization priors, summarized in Figure 5 D as normalized utility, model-derived spike probability, and entropy across HPRs for each group. Among all-unit HHL contrasts, directional support was greatest at the quietest context (HPR 30), where NE shows higher normalized utility than SH (median Δ*Ū* = − 0.23, 95% CI [−0.53, 0.04], Pr(Δ *<* 0) = 0.95; Table 2), despite lower implied firing and lower entropy in the matched priors. NE priors were also more concentrated (lower entropy) than SH at quiet contexts (Pr = 0.96 at HPR 30; Pr = 0.79 at HPR 45; Table 2), consistent with the empirically confirmed reduction in gain modulation. In this context, entropy describes the spread of the inferred prior over threshold-gain configurations. A firing-rate crossover accompanied the utility pattern: SH showed higher implied firing at quiet (Pr = 0.95 at HPR 30) while NE showed higher implied firing at the loudest context (Pr = 0.82), consistent with gain headroom exhaustion limiting the ability of noise-exposed neurons to suppress activity at loud levels. Transition contrasts further indicated that the NE quiet-context advantage eroded progressively as the acoustic environment became louder (Pr = 0.91-0.96 for transitions from HPR 30 to louder contexts; Table 2). Together, these directional patterns (utility, entropy, firing rate, and transitions) are consistent with a quiet-context advantage that depends on how prior mass is organized over threshold-gain space rather than on a simple increase in activity. Notably, this quiet-context separation was not mirrored by threshold modulation and was not cleanly resolved by Fisher information at the animal level (Table 1), indicating that the optimization-prior analysis captures aspects of the information-cost structure that local sensitivity measures alone do not separate. At higher HPRs, the HHL utility contrast becomes weaker and more heterogeneous: the advantage at HPR 45 narrows substantially, HPR 60 behaves as a transitional regime, HPR 75 shows only small utility differences in a lower-*ξ* state, and at the loudest context the NE trajectory approaches the *ξ* = 0 boundary while both groups exhibit further utility loss and sharply increased implied firing.

**Table 2.** Table S2. Optimization-prior inference contrasts reported in the main text (balanced bootstrap, 3 000 replicates). Ranked by directional support Pr_dir_, the proportion of bootstrap replicates consistent with the predicted direction. Markers are assigned from unrounded probabilities: ** for strong support (Pr_dir_ ≥ 0.95), * for moderate (0.85 ≤ Pr_dir_ *<* 0.95); unmarked rows are inconclusive per the evidence categories defined in Section. CHL OP contrasts (all HPRs, all quantities) yielded Pr_dir_ *<* 0.73 and are omitted for brevity.

| ID | Contrast | HPR | Population | Median bootstrap $\Delta$ | 95% CI | Directional support $Pr_{dir}$ |
| --- | --- | --- | --- | --- | --- | --- |
| TX1 | Utility transition 30→45 | 30→45 | All units | +0.21 | [-0.03, +0.45] | 0.961** |
| HH1 | Entropy (SH - NE) | 30 | All units | +0.24 | [-0.02, +0.50] | 0.959** |
| HE1 | Utility (SH - NE) | 30 | Low-threshold | -0.22 | [-0.51, +0.03] | 0.956** |
| TX2 | Utility transition 30→60 | 30→60 | All units | +0.29 | [-0.05, +0.63] | 0.953** |
| HD1 | Utility (SH - NE) | 30 | All units | -0.23 | [-0.53, +0.04] | 0.953** |
| HF1 | Mean FR (SH - NE) | 30 | All units | +0.09 | [-0.02, +0.23] | 0.946* |
| TX3 | Utility transition 30→75 | 30→75 | All units | +0.25 | [-0.05, +0.60] | 0.942* |
| TX4 | Utility transition 30→90 | 30→90 | All units | +0.21 | [-0.09, +0.58] | 0.909* |
| S5 | Utility (SH - NE) | 90 | High-threshold | +0.14 | [-0.09, +0.39] | 0.873* |
| HF5 | Mean FR (SH - NE) | 90 | All units | -0.04 | [-0.12, +0.00] | 0.824 |
| HH2 | Entropy (SH - NE) | 45 | All units | +0.09 | [-0.13, +0.32] | 0.787 |
| HE5 | Degree of divergence | 60 | Stratified | +0.05 | [-0.21, +0.30] | 0.644 |
| S1 | Utility (SH - NE) | 30 | High-threshold | -0.05 | [-0.29, +0.14] | 0.640 |

#### A two-population decomposition shows the quiet-context HHL effect distributed across low- and intermediate-threshold neurons

Because HHL selectively targets high-threshold, low spontaneous-rate auditory-nerve fibers, the all-unit optimization-prior contrasts may reflect a mixture of neurons differentially affected by the lesion. We therefore hypothesized that threshold-defined subpopulations would carry unequal shares of the quiet-context utility shift, with the high-threshold population (whose afferent input is most affected by the lesion) showing an attenuated NE advantage consistent with deafferentation.

To test whether the all-unit HHL contrast reflects a uniform shift of the entire population or a change concentrated in a subset of neurons, we split the fitted threshold-gain samples into low- and high-threshold populations and repeated the same optimization-prior inference (Figure 5 E). Neurons were assigned to low- and high-threshold groups by their median adapted threshold across contexts, split at 85 dB SPL (the upper third of the tested 24-96 dB SPL range, which also coincides with the boundary at which directional NE > SH support collapses across HPRs). This decomposition uses the same SH-NE utility contrast as the all-unit analysis but determines whether quiet-context differences are carried preferentially by one threshold-defined subpopulation.

The quiet-context NE advantage is localized to the low-threshold population (low-threshold HE1, 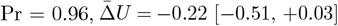; Table 2), whereas the high-threshold population shows no group difference at HPR 30 (high-threshold S1, Pr = 0.64, 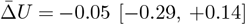) and reverses direction at the loudest context (Pr(SH *>* NE) = 0.87, 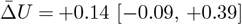 at HPR 90; Figure 5 E).

Resolving the contrast across the full range of adapted thresholds (Table 3) refines this pattern. At quiet contexts (HPR 30 and 45) the NE advantage spans low- and intermediate-threshold neurons (Q2 at HPR 45, 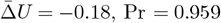; Q3 at HPR 30,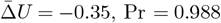; Table 3) and is lost—or reverses toward SH—only in the highest-threshold neurons (≥ 89 dB SPL; Pr(NE *>* SH) = 0.52 at HPR 30 and 0.27 at HPR 45; Table 3). This loss specifically in the highest-threshold neurons is consistent with the selective deafferentation of high-threshold afferent fibers that characterizes hidden hearing loss. At louder contexts the threshold profile is weak and heterogeneous.

**Table 3.** Table S3. Threshold-resolved directional support for the HHL utility contrast. Proportion of bootstrap replicates with a quiet-context NE advantage, Pr(NE *>* SH) on normalized utility, for neurons grouped into equal-sized quintiles of across-context median adapted threshold. Values *<* 0.5 indicate a reversal toward SH. Markers are assigned from the displayed probabilities: ** for Pr ≥ 0.95, *** for Pr ≥ 0.99, matching the convention in Figure 5 E. At quiet contexts (HPR 30 and 45) the advantage spans low- and intermediate-threshold neurons and falls off in the highest-threshold quintile; louder contexts are heterogeneous.

| Adapted threshold (dB SPL) | $n$ (NE/SH) | HPR 30 | HPR 45 | HPR 60 | HPR 75 | HPR 90 |
| --- | --- | --- | --- | --- | --- | --- |
| 24-54 | 92/144 | 0.878 | 0.631 | 0.578 | 0.390 | 0.557 |
| 54-61 | 127/108 | 0.949 | 0.959** | 0.258 | 0.764 | 0.835 |
| 61-72 | 118/117 | 0.988** | 0.906 | 0.753 | 0.914 | 0.298 |
| 72-89 | 88/147 | 0.914 | 0.831 | 0.858 | 0.759 | 0.738 |
| 89-96 | 125/111 | 0.517 | 0.272 | 0.364 | 0.253 | 0.119 |

This threshold dependence is consistent with peripheral computational models of fiber-type contributions (Johannesen et al., 2022). Among the all-unit per-HPR contrasts, no degree-of-divergence effect emerges at HPR 60 under the 85 dB SPL split (Pr = 0.64; Table 2), consistent with the broader pattern that HPR 60 behaves as a transitional regime rather than a stable locus of group separation. We interpret the quiet-context HHL effect as distributed across low- and intermediate-threshold neurons and attenuated in those with the highest adapted thresholds (Table 3), rather than a uniform shift or a strict low-threshold locus.

At HPR 90, the posterior frequently concentrates near the boundary *ξ* → 0 (occurring in more than 85% of NE bootstrap samples), so we interpret this condition cautiously rather than as a stable group-comparison point. In this limit regime, once the loudest context drives the system into a high-activity state where the firing-rate penalty is effectively irrelevant, the prior loses organization and the model no longer cleanly discriminates among lesion conditions.

For CHL, plug removal (PP) shifts the inferred priors toward slightly greater organization and utility than the plugged state (EP), but these differences are modest and considerably weaker than the main HHL quiet-context effect. Model-derived spike probability remains broadly similar between EP and PP over 30-60 dB SPL, whereas entropy is modestly lower after plug removal. We therefore treat the CHL optimization-prior results as supportive evidence for a distinct lesion trajectory rather than as a stand-alone strong contrast.

Together, the trajectories and their associated statistics indicate a general operating principle: as mean intensity increases, the system first relaxes the firing-rate penalty (lower *ξ*) while tightening the prior around highly informative threshold-gain configurations (higher *β*), but at sufficiently loud contexts the cost constraint saturates and the inferred code enters a diffuse, high-activity regime. Pathological conditions may therefore be better distinguished by their context-dependent trajectories, implied firing, and organizational structure than by any single HPR-local sensitivity measure.

## Discussion

Sensory systems must continually adapt their neural representations to match the prevailing statistics of the environment, reallocating limited dynamic range to encode the most informative regions of stimulus space. When sensory input is perturbed by subtle lesions or changes in input gain, these fast-adapting mechanisms can be pushed into operating regimes that preserve basic sensitivity yet alter how information is distributed across neurons and contexts. In this sense, hidden hearing loss (HHL) is an instance of a subtle lesion that currently poses a clinical paradox: substantial listening difficulty in noise despite clinically normal thresholds. Here we put HHL (in the form of noise-induced synaptopathy) and experimentally induced conductive attenuation (ear plugging) and its removal, within a single, quantitative framework that evaluates how neural operating points move across different listening environments (HPR contexts). This interpretation is anchored by the confirmed empirical finding that gain modulation across acoustic contexts is significantly altered in noise-exposed animals with otherwise normal audiometric thresholds (group × context interaction, *p* = 0.009, corrected), establishing a physiological substrate for the optimization-prior trajectory differences. Further, rather than exploring the consequences of HHL and CHL in isolation, we use the same optimization-prior landscape to determine how, at a systems level, the auditory brain reconfigures its coding parameters in response to changes in the sensory environment (mean background intensity). This allows us to make direct contrasts between pathologies using the same coordinates and the same stimulus parameters. From this, we suggest these dynamics might be linked to changes (deficits, but not only) in performance in listening tasks for these two forms of hearing impairment.

### A single normative landscape captures modified neural coding in response to subtle sensory lesions

A key goal of our approach is to generate a single framework under which changes in sensory input might be understood in terms of how neural systems allocate their resources to accommodate a wide range of stimulus possibilities, including when peripheral sensory processing is altered. Here, we modeled adaptation to statistically defined distributions of sound intensities as movement on a two-parameter landscape defined by optimization priors over threshold-gain configurations. The priors were constructed from a utility function trading mutual information about stimulus categories against a metabolic penalty, namely mean firing rate. We found that fitting a family of priors to neural responses across different sound environments defined by their statistical distribution of sound intensities yielded trajectories in the information-metabolism plane (*β, ξ*) that summarize how the system reallocates dynamic range as the mean of the distribution of sound intensities increases. Gjorgjieva et al. (2019) et al. recently demonstrated that, with the objective of efficient coding, the distribution of thresholds can be predicted from stimulus statistics and constraints of intrinsic noise and metabolic cost, and that different objective functions (e.g., MI *vs*. error-based criteria) can yield different apparent “optimal” allocations. This interpretation is supported by Röth et al. highlighting that the source of noise and downstream neural convergence (pooling of multiple afferent channels onto single neurons) lead to a reorganization of the distribution of neural thresholds as effective noise increases (Röth et al., 2021). Recent work has further extended this class of normative models by jointly optimizing encoding and decoding, linking efficient sensory representations to learnable generative objectives (Blanco Malerba et al., 2024). As such, the pathology-dependent (*β, ξ*) trajectories we infer can be interpreted as lesion-driven shifts in effective constraints (cost pressure and/or effective noise) that modulate whether, and the extent to which, neural parameters concentrate near high-utility regions, even when classic sensitivity metrics within the HPR appear similar. Across groups, the 30→45 transition relaxes the rate constraint (lower *ξ*) whilst increasing information/organizational pressure (higher *β*), the 45→60 transition partially reverses this tendency, and trajectories return toward lower-*ξ* states at higher HPRs; at the loudest context, boundary behavior dominates and normalized utility declines despite high firing. Research in other modalities has shown that related efficient-coding principles can account for central visual sensitivity to higher-order texture statistics (Tkačik et al., 2010) and for structured interactions within hippocampal place-cell networks, indicating that information-metabolism trade-offs are not confined to the sensory periphery. Considered in this context, the auditory optimality landscape we derive here can be seen as one instance of a more general principle: using the same normative coordinates to compare how different sensory modalities and circuit levels redistribute limited resources when the structure or reliability of their inputs changes.

### A continuum of coding regimes from CHL to HHL

In the information-metabolic plane (*β, ξ*), conductive attenuation of sound produces a distinct envelope, with ear-plugged and post-plug states occupying lower-*β*, higher-*ξ* regions than HHL conditions (SH and NE groups). This CHL envelope remains largely confined to an iso-contour corridor characteristic of a compressed dynamic range and terminates near the region where HHL control trajectories begin.

Compared to neural responses from control animals with no known pathology, responses of neurons in noise-exposed (NE) animals with evidence of HHL showed larger reductions in the metabolic penalty (*ξ*) as the mean intensity of a sound environment increased, but only modest increases in *β* between quiet and moderately loud sound environments. This yields the clearest utility advantage in quiet conditions, despite lower prior entropy and lower implied firing in the matched priors at HPR 30 (a pattern consistent with cortical evidence that cochlear neural degeneration disrupts coding in background noise (Resnik and Polley, 2021) and with models linking compensatory gain changes to perceptual deficits (Johannesen and Lopez-Poveda, 2021; Zeng, 2020)). Our two-population analysis indicates that this quiet-context effect spans low- and intermediate-threshold IC neurons and is attenuated in those with the highest adapted thresholds (Table 3). This complements mechanistic HHL models that distinguished synaptopathy from myelin disruption using forward models of auditory-nerve activity (Budak et al., 2021) by locating lesion-dependent IC adaptations on common information-cost axes. In contrast, responses of neurons in SH controls (no known HHL or CHL pathology) maintain higher organization (*β*) at comparable rate/metabolic penalty (*ξ*), indicative of more organized priors and lower rates across different listening contexts.

CHL occupies a different and weaker region of the same landscape. EP and PP responses remain shifted toward lower *β* and higher *ξ* than the HHL conditions, and plug removal produces only a modest shift toward greater organization and utility (canonical CHL optimization-prior contrasts remained inconclusive, Pr_dir_ *<* 0.73). This outcome must be considered in the context of the short delay between EP and PP recordings; a longer interval between recordings may have revealed changes to these parameters arising over a longer time course. Nevertheless, this pattern is consistent with incomplete rapid renormalization after conductive attenuation rather than with a large reconfiguration of coding strategy over the timescale measured here.

Importantly, across animal groups and conditions, the representation in the neural optimality landscape converges at loud contexts to an unrestrained rate/firing saturation regime (low *ξ*) where utility advantages collapse, even as differences in organizational structure and implied rate persist. At the loudest context, this regime becomes boundary-dominated, especially for NE, so the framework is most informative for quiet-to-moderate contexts and treats the loudest context as a limit case rather than a clean inferential comparison. Because all manipulations are interpreted in the same coordinate system and over the same HPR transitions, these distinctions constitute a differential, mechanistic account rather than a collection of case-specific phenomena.

### From energetic sensitivity to informative assessment

Traditional energetic measures (thresholds, average rates, or even Fisher information confined to an HPR) can obscure meaningful system-level differences that emerge as stimulus distributions unfold over multiple timescales and are likely to impact perception across tasks. Indeed, none of the group comparisons of FI in our data survived correction for between-animal variance in linear mixed-effects models (Table 1), underscoring the importance of the optimization-prior framework for detecting system-level differences that FI alone cannot resolve. Two populations can show similar sensitivity within an HPR while differing markedly in how much spiking they expend outside that region and in how tightly they concentrate probability mass on informative threshold-gain configurations. By quantifying an explicit trade-off between information and metabolic cost, the optimization-prior landscape provides candidate bridge variables (utility, entropy, and implied firing rate) between spike trains and psychoacoustic performance, shifting the emphasis from energetic sensitivity to informative assessment. The improved normalized utility observed in quiet environments for noise-exposed animals is thus reconciled with an overall reduction in efficiency once organizational spread and rate expenditure are accounted for, and the two-population analysis shows this quiet-context effect is distributed across low- and intermediate-threshold neurons and attenuated at the highest adapted thresholds rather than reflecting a uniform shift of the entire HHL ensemble, whereas conductive attenuation maps to information-poor, energy-frugal regimes that shift only modestly with context. In this view, behavioral limits are expected to depend not only on local sensitivity but also on how reliably and economically information is distributed across the full stimulus distribution that a listener experiences.

Our approach is related to recent work in which deep neural networks trained on ecologically relevant auditory tasks reproduce aspects of human-like perception (Saddler and McDermott, 2024; Francl and McDermott, 2022), and to population encoding models that generalize across neural populations in auditory cortex (Pennington and David, 2023), reinforcing the role of stimulus statistics in shaping neural coding. By distilling population-level coding into a two-parameter information-cost trade-off, the optimization-prior framework offers an interpretable, low-dimensional alternative that is naturally suited to detecting the subtle system-level shifts that high-dimensional models may not resolve.

### Translational implications and testable predictions

As the same landscape coherently characterizes synaptopathy and conductive attenuation, this can guide stimulus selection toward contexts and transitions that maximize separability of trajectories across that landscape, yielding more informative assessments than relatively simple measures (e.g. sound-evoked neural thresholds). Closed-loop paradigms that adjust mean intensity to probe iso-contour boundaries or the onset of low-*ξ* saturation, for example, are natural extensions of this approach and can, in principle, be implemented with population electrophysiology. With that aim in mind, several testable hypotheses follow directly from the trajectory geometry. Following plug removal, for example, re-exposure to speech or natural soundscapes should progressively increase organization (*β*) and allow the rate penalty (*ξ*) to relax in a time-dependent fashion, reducing prior entropy across contexts. Progressive synaptic compromise should generate persistent down-shifts in *ξ* with attenuated gains in *β* from quiet to moderate contexts, shrinking any quiet-context advantage, whilst pharmacological manipulations that weaken inhibition should lower *ξ* and, if instability ensues, reduce *β*. Conversely, strengthening inhibition should have the opposite effect, moving trajectories toward higher organization at fixed utility. Analogous trajectory families should be observable in other modalities under distributional stimulation, with modality-specific mappings between circuit constraints and (*β, ξ*).

Our current account is deliberately phenomenological, prioritizing interpretability of operating regimes over biophysical detail. We estimated priors on a finite threshold-gain grid using sigmoidal response families and a discrete mutual information estimator with category binning relative to the HPR, under data-rich and multiple-instance simplifications and with hierarchical bootstrap resampling at the animal level. These choices bring tractability and transparent geometry to the problem but also delimit scope: anesthesia, single-site recordings, binarized responses, finite grids, and estimator bias all constrain absolute values and may blur fine-grained differences. All recordings were obtained under ketamine/xylazine anesthesia; because the same protocol was used for all groups, between-group contrasts remain internally valid, but absolute hyperparameter values should be treated as relative, and awake recordings would be needed to confirm that the trajectory geometry generalizes beyond the anesthetized preparation. The available sample sizes remain modest (4 NE, 6 SH after SH1 exclusion for HHL LME tests, 4 EP, and 4 PP animals), unit yields are uneven across animals, and the HHL threshold-gain tables include repeated fit variants for some units, so contrasts across HPRs and subpopulations are best interpreted as directional and mechanistic rather than as a family of independently confirmed tests. We note that the multiple comparisons across HPR levels, lesion types, and subpopulations were not subjected to formal family-wise error correction; the bootstrap-derived directional-support statistics are therefore best read as posterior summaries of effect direction and magnitude rather than as classical hypothesis tests. The coherent pattern of multiple moderate-evidence contrasts (utility, entropy, firing rate, and transitions all pointing in the predicted direction at quiet contexts) is more informative than any single contrast in isolation, but this convergence should be confirmed in adequately powered replication studies. Inference at loud contexts is additionally sensitive to boundary behavior, especially when *ξ* approaches zero, which is why we emphasize quiet-to-moderate contexts and treat the loudest context as a limit case of the framework rather than a stable group-comparison point. None of these caveats undermines the qualitative claim that subtle lesions occupy distinct envelopes in the same normative coordinates and that adaptation can be read as trajectories on that landscape, but they do set the evidentiary bar for translational use. Longitudinal designs will be particularly informative: tracking recovery from conductive hearing loss or the evolution of noise-induced changes in hearing and listening performance over weeks to months should reveal predictable realignments of *β* and *ξ* across contexts, directly testing the stability and specificity of trajectory features. Extending beyond the auditory midbrain to multi-area recordings, relaxing the binary-response assumption, and incorporating sparsity or alternative coding costs will increase the reach of the framework while preserving its central interpretive advantage: a single, rigorous coordinate system in which distinct subtle lesions and their adaptations can be compared across contexts and over time. Translational versions should be tied to established human-compatible HHL biomarker approaches, including ABR latency-in-noise and envelope-following-response measures (Valderrama et al., 2022; Mehraei et al., 2016; Shaheen et al., 2015).

## Materials and methods

### Datasets

The HHL dataset consists of additional recordings obtained in the same animals and experimental sessions as those reported in (Monaghan et al., 2020). The CHL dataset was collected in the same laboratory using the same recording setup and analysis procedures.

All experiments were conducted under a United Kingdom Home Office project license (30/2481) and complied with the 1986 Animals (Scientific Procedures) Act. Adult male Mongolian gerbils (Meriones unguiculatus) were used, weighing 70-90 g and aged 3-6 months. Animals were randomly assigned to one of four experimental groups: sham-exposed, noise-exposed, ear-plugged, or post-plug. Total numbers of animals were 4 noise-exposed (NE), 6 sham-exposed (SH), 4 ear-plugged (EP), and 4 post-plug (PP) gerbils; the EP and PP groups represent the same 4 physical animals recorded before and after plug removal. For empirical linear mixed-effects models, analyses used animals with sufficient data for the corresponding measure; animal counts are reported with each statistic. For the optimization-prior inference, all 6 SH animals contributed to the bootstrap.

### Protocols timeline

For the HHL condition, auditory brainstem responses (ABRs) were first measured on day 1 to determine baseline hearing thresholds in all animals. ABR methods and example traces are described in detail in Monaghan et al. (2020) and are not repeated here. On day 2, animals assigned to the noise-exposed group were subjected to a high-intensity noise exposure (see below), while sham animals underwent anesthesia only. On day 3, ABRs were measured again to confirm a temporary elevation in hearing thresholds caused by the noise exposure. On day 30, ABRs were measured a third time to verify the recovery of thresholds. Immediately after these recordings, we made extracellular recordings from neurons in the inferior colliculus (IC) using multi-electrode arrays.

For the CHL condition, on day 1, animals were anesthetized and fitted bilaterally with foam earplugs covered with otoform impression material. ABRs were measured immediately before and after earplug insertion to quantify the induced conductive loss. On day 8, ABRs were recorded again, followed by extracellular IC recordings with the earplugs still in place. At the end of this recording session, the earplugs were removed and IC activity was recorded again immediately to obtain the post-plug (PP) condition.

### Noise exposure in the HHL group

For noise exposure, gerbils were anesthetized with an intraperitoneal injection of a fentanyl/medetomidine/midazolam mixture (same drug ratios as in the recording procedures described below). Supplemental doses were given as needed based on the pedal withdrawal reflex. Body temperature was maintained at 37-38 °C and respiration was monitored every 30 minutes.

Animals were placed in a custom-built sound-attenuating chamber under a loudspeaker (Monacor Stage Line PA Horn Tweeter MHD-220N/RD, Bremen, Germany) positioned 45 cm above the animal’s head. Before each experiment, the loudspeaker output was calibrated to be flat (±2 dB) between 2 and 4 kHz at the position of the animal. Noise-exposed animals were then exposed to 2-4 kHz octave-band noise at 105 dB SPL for 2 hours, generated and controlled using TDT System 3 hardware. After the noise exposure, animals received an intraperitoneal injection of atipamezole/flumazenil/naloxone (same ratios as above) to reverse anesthesia and were allowed to recover for about 1 hour in a temperature-controlled chamber. Sham-exposed animals underwent the same procedure except that the loudspeaker remained silent.

### Single-neuron recordings

Extracellular recordings were made in a sound-insulated booth (Industrial Acoustics, Winchester, UK). Anesthesia was induced with an intraperitoneal injection of a solution containing ketamine (100 mg/mL), xylazine (2% w/v), and saline mixed in a 5:1:19 ratio as described in Schnupp et al. (2015). The same anesthetic mixture was infused continuously throughout the experiment at approximately 2.1 *µ*L/min via an intraperitoneal line. Body temperature was held at 38.7 °C with a feedback-controlled heating blanket.

After making a midline scalp incision, the skull was exposed and a small metal head pin was cemented to the skull. The head pin was fixed to a stainless-steel head holder in a stereotaxic frame. A craniotomy was then made over the right IC, extending about 3.5 mm from the midline and centered on the lambdoid suture.

Neural activity was recorded using 64-channel silicon probes (Neuronexus Technologies, Ann Arbor, MI, USA) arranged as tetrodes, with two tetrodes per shank and four shanks per probe. After removing the dura, the probe was advanced through the overlying cortex. To minimize tissue damage and drift, the probe was first advanced at 1 *µ*m/s and slowed to 0.3 *µ*m/s as it approached the IC. Recording sites were positioned within the central nucleus of the IC, identified by the presence of tonotopically ordered frequency tuning across the array. Once responses to the full set of stimuli had been collected at a given depth, the probe was advanced by approximately 300 *µ*m and another recording run was obtained. Typically, at least three penetrations were made along the rostro-caudal axis at 150-*µ*m intervals, and data were collected at two depths in each penetration. During recordings, ECG and core body temperature were monitored continuously, and oxygen-enriched air was delivered near the snout.

Spike sorting followed the procedures described in Garcia-Lazaro et al. (2015). In brief, signals from each channel were bandpass filtered between 500 and 5000 Hz and then whitened on a tetrode-by-tetrode basis to decorrelate the four channels. Candidate spikes were detected as waveform snippets whose energy (Choi et al., 2010) exceeded a predefined threshold, with a minimum inter-spike interval of 0.7 ms. Each snippet was projected onto the first three principal components for each channel, and spike clusters were identified in this feature space using KlustaKwik (http://klustakwik.sourceforge.net) and Klusters (Hazan et al., 2006). The quality of each cluster was quantified using the isolation distance metric described in Schmitzer-Torbert et al. (2005), which assumes that clusters form multivariate Gaussian clouds in feature space and measures how far the cluster must expand (in SD units) to double the number of included spikes. For each tetrode, the “noise” or multiunit cluster always contained at least as many spikes as any single-unit cluster (Garcia-Lazaro et al., 2015). For the main optimization-prior analysis, we used the HHL and CHL datasets described above and retained units whose rate-intensity functions were fit by the sigmoid model with mean squared error <= 0.08, without excluding units by gain. Uncertainty for all main-text estimates was quantified with hierarchical bootstrap resampling at the animal level (45 threshold-gain samples per replicate for HHL and 30 for CHL). Because some HHL units contributed more than one threshold-gain fit variant, we report animal counts explicitly and show per-animal sample composition in Figure 5 E rather than quoting a single pooled single-unit total. Within each HPR condition, each neuron contributes exactly one threshold-gain pair; a neuron contributes separate fits across HPR conditions, which is appropriate because each HPR represents a distinct statistical context producing a different adapted response.

### Empirical group comparisons

We report animal-level summaries for all empirical claims (threshold shifts, spike-rate differences, tuning-curve statistics). For each measure, we first computed the per-animal mean, then compared groups with a Welch two-sample *t*-test. Effect sizes are given as Hedges’ *g* with bias-corrected 95% confidence intervals.

For tests of context-dependent effects (group × HPR interactions), we fitted linear mixed-effects models (LMEs) with group, HPR (categorical), and their interaction as fixed effects and animal as a random intercept, using restricted maximum likelihood estimation. The joint significance of the group × HPR interaction was assessed with a Wald *χ*^2^ test on the interaction coefficients. For main-effect-only comparisons, LMEs with group as the sole fixed effect and animal as random intercept were used. Where multiple claims are tested within the same family, *p*-values are adjusted with the Benjamini-Hochberg false-discovery-rate procedure. Five families were defined to reflect independent scientific questions: HHL-core (gain modulation, threshold modulation, spontaneous rate), Fisher information (total FI, localized FI, dynamic range), distributional spread (gain spread, threshold spread), CHL within-subject (threshold, rate, gain modulation, FI, spontaneous rate), and cross-cohort (EP vs SH, EP vs NE, PP vs SH). Pooled single-neuron analyses (treating every unit as independent) are reported as *descriptive* supplements; because units within an animal share surgical, acoustic, and physiological conditions, pooled tests inflate effective sample size and are not used for inferential conclusions. Full results appear in Table 1.

### Optimization-prior inference

Optimization-prior (OP) comparisons quantify *directional support* for a hypothesis rather than classical significance. For each hypothesis we compute a bootstrap distribution of the relevant contrast (e.g., ΔMI between NE and SH) using a hierarchical bootstrap that resamples animals, then neurons within animals, for 3,000 iterations. We then report: (i) the median contrast and its bootstrap 95% CI (2.5th-97.5th percentile); (ii) the directional probability Pr(Δ < 0) or Pr(Δ > 0), whichever is the predicted direction; and (iii) an evidence label: *strong* (Pr ≥ 0.95), *moderate* (0.85 ≤ Pr < 0.95), or *inconclusive* (Pr < 0.85). These thresholds parallel the Bayesian evidence categories of Kass and Raftery (1995): directional odds of approximately 6:1 (moderate) and 20:1 (strong), adopted conservatively given the available sample sizes. Standardized effect sizes (bootstrap Cohen’s *d* with 95% CI) are reported alongside directional probabilities in Table 2.

No classical *p*-values or significance thresholds are applied to optimization-prior quantities.

All HPR levels are retained in inferential summaries; boundary behavior at the loudest context is interpreted cautiously in Results.

Optimization-prior results reported in the main text are given in Table 2.

### Stimuli

To investigate auditory intensity coding capacity, we used a stimulus that consists of broadband noise (2-45 kHz) whose intensity was changed every 50 ms, each intensity being a realization of a distribution with a high probability region (HPR), depicted in Figure 1 A, D. The range of intensities was always integer, between 24 and 96 dB SPL, with 3 dB steps. In total, a sequence of 2000 intensities (realizations) were presented per HPR-distribution, resulting in recordings of 100 s for each HPR-distribution.

### Posterior distribution calculation

Let 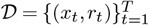 be a dataset composed of *T* stimulus-response pairs, coming from a single neuron (or a population of them), that can be represented with parameters *θ* = {*x*_0_, *k*}, to be estimated. The posterior over *β* takes the form

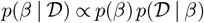

where *p*(*β*) is the prior distribution of *β*, here regarded as uniform, and the likelihood *p*(𝒟 | *β*) defined as

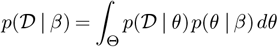

In this equation, *p*(𝒟 | *θ*) represents the marginal likelihood of neuronal parameters *θ*, estimated from dataset 𝒟, and *p*(*θ* | *β*) the optimisation prior distribution, derived from a normative theory (efficient coding in this case). In this way, the magnitude of the posterior *p*(*β* | *𝒟*) depends on the overlap between the likelihood of neural parameters *p*(𝒟 | *β*) with the optimisation prior distribution *p*(*θ* | *β*). An estimate 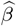 is yielded via MAP, chosen as the value of *β* that maximises its posterior distribution, i.e.

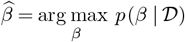

#### Mathematical simplifications

##### Data-rich regime simplification

If the dataset 𝒟 has sufficient samples to estimate accurately the value of *θ*, the likelihood *p*(𝒟 | *θ*) will carry most weight around the estimated parameters 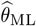, namely the parameters with maximum likelihood (ML). Thus, in this case, the magnitude of the posterior *p*(*β*| 𝒟) will depend on the value of the optimisation prior at that point, i.e.,

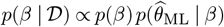

Equally, the estimations of parameters 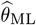 in this data-rich regime can be achieved using any function-fitting algorithm, with the aim of minimising the mean squared error.

##### Multiple system instances simplification

The equations derived above consider only a single neuron (or a population of them), represented by a single parameter *θ*. The following extension enables the assessment of a population of neurons, represented through a set of parameters {*θ*_1_, …, *θ*_*N*_}, that are assumed to arise from an optimisation prior distribution *p*(*θ* | *β*), for a *β* to be inferred via MAP.

Let {𝒟_1_, …, 𝒟_*N*_} be a set of *N* datasets, each with corresponding parameters 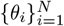, respectively. Assuming that the datasets are i.i.d., the parameters 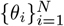 are also i.i.d., so the joint probability distributions can be expressed as,

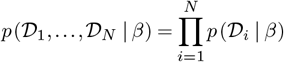

this simplification renders the posterior as

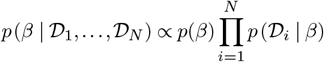

Putting together both simplifications, the magnitude of the posterior distribution over *β* can be estimated as

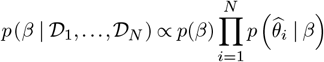

where 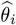 corresponds to the estimate of neural parameters via maximum likelihood estimation or minimisation of mean squared error. MAP uncertainty was quantified with 3000 hierarchical animal-bootstrap replicates

##### Rate-intensity functions

Input/output functions (referred to here as rate-intensity functions, RIFs) were estimated for each single-unit by averaging the spike count per intensity. An example of a single-unit RIF is shown in Figure 1 D, for all HPR-distributions, centered at 30, 45, 60, 75, and 90 dB SPL and no-HPR (uniform). The solid line is the fit to the RIF of a logistic function of the form

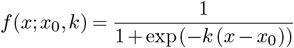

where *x* is the intensity axis, *x*_0_ the threshold and *k* the gain or slope of the sigmoid. These last two parameters {*x*_0_, *k*} are plotted as a bow tie symbol in Figure 1 D; the black line refers to the uniform intensity distribution.

All single-unit RIFs were averaged in order to estimate a population response, shown in the upper panel of Figure 2 and Figure 3. Fisher information (FI) was estimated from a spline-smoothed version of the population RIF, then applying a 6 dB Gaussian filter along the intensity axis, shown in the second row of Figure 2 and Figure 3. To obtain the FI as a function of the intensity *x*, the following approximation was employed (Harper and McAlpine, 2004),

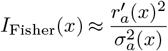

where *r*_*a*_(*x*) is the population (average) RIF and 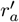 its derivative; 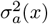 is the variance of *r*_*a*_ for each intensity, which we approximated using a Poisson assumption over *r*_*a*_ *x*), thus 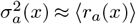.

Single-unit RIFs were fitted to logistic functions of the form

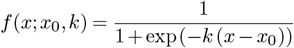

where *x* is the intensity axis, *x*_0_ is the threshold and *k* is the gain. A two-dimensional parameter grid for *x*_0_ and *k* was built as a 25 × 25 matrix, using 25 levels for *x*_0_ and *k*, with *x*_0_ ∈ [24, 96] and *k* ∈ [− 0.5, 0.5]. Fits were obtained by evaluating this discrete grid directly after response normalization and selecting the parameter pair that minimized mean squared error. All RIFs were first normalized in order to estimate only parameters {*x*_0_, *k*}, setting the spontaneous rate to 0 and the maximum rate to 1.

## Notes

### Competing Interest Statement

The authors have declared no competing interest.

### Summary of Updates

Supplementary tables updated, added funding information.

### Funding

- Australian Research Council: DP180102524

